# Nighttime Caffeine Intake Increases Motor Impulsivity

**DOI:** 10.1101/2025.06.09.658656

**Authors:** Erick Benjamin Saldes, Paul Rafael Sabandal, Kyung-An Han

## Abstract

Caffeine is commonly consumed at night by shift workers and military personnel for its wake-promoting effect, yet its adverse effects on behavior remain underexplored. Here, we show that nighttime caffeine intake impairs inhibitory control in *Drosophila melanogaster*, resulting in impulsive motor behavior, with females more sensitive than males. This effect is independent of hyperactivity or sleep loss, as walking speed was unchanged, and artificial sleep deprivation via light or mechanical stimulation did not elicit similar deficits. Notably, daytime caffeine feeding did not impair inhibitory control, highlighting a circadian gating of caffeine’s behavioral impact. Mechanistically, we identify dopamine signaling as a key mediator of caffeine-induced impulsivity. Reduced dopamine synthesis (*pale/+*), silencing of PAM dopaminergic neurons, or altered dopamine transporter activity (*fumin/+*) attenuated or exacerbated caffeine-induced impulsivity. Targeted manipulations identified the dopamine D1 receptor (dDA1/Dop1R1) in the mushroom body (MB) α/β and γ lobes as essential for this effect, with γ-lobe neurons exhibiting heightened sensitivity. These findings reveal a circadian- and dopamine-dependent mechanism through which nighttime caffeine impairs behavioral inhibition.

## Introduction

Caffeine is the most widely consumed psychoactive substance in the world, with approximately 85% of adults in the United States reporting regular use^1^. Its popularity stems from its ability to increase alertness, reduce fatigue, and improve performance, especially under conditions of sleep deprivation. For these reasons, caffeine is routinely used by shift workers, healthcare professionals, military personnel, and others who must remain vigilant during nighttime hours^2–8^. A growing body of research supports caffeine’s effectiveness in enhancing reaction times, vigilance, and attention in high-stress or sleep-deprived settings^4,9–12^.

Despite its benefits, caffeine consumption is also associated with a range of adverse effects. At higher doses or in caffeine-sensitive individuals, it can induce anxiety, restlessness, nausea, jitteriness, and trembling^13–15^. Several studies have also reported that caffeine can impair fine motor control. For example, caffeine-naïve individuals have been shown to exhibit reduced hand steadiness, decreased manual dexterity, and increased performance errors following even moderate caffeine consumption^15^. These impairments may compromise motor accuracy and increase the risk of accidents or errors in occupational settings where precision is critical.

Although caffeine’s physiological and behavioral effects are well established, the underlying mechanisms, particularly those contributing to caffeine-induced impairments in cognitive and motor functions, remain poorly understood. Furthermore, while most studies have focused on caffeine’s effects during daytime or on sleep loss per se, its behavioral impact following nighttime consumption remains largely unexplored, despite its widespread use among overnight shift workers. To address these knowledge gaps, we used *Drosophila melanogaster* as a model system to investigate how nighttime caffeine intake affects inhibitory control, a fundamental executive function responsible for suppressing inappropriate actions. *Drosophila* serves as a powerful genetic model for studying conserved mechanisms of neurotransmission, synaptic plasticity, and high-order functions such as learning and memory across species.

In this study, we show that nighttime caffeine consumption in *Drosophila* impairs inhibitory control and induces motor impulsivity. This effect is sexually dimorphic, independent of hyperactivity or sleep loss, and mediated by dopamine signaling within the mushroom bodies, a neural structure involved in high-order functions such as learning, memory, and decision-making^16–19^. Our findings uncover a previously unrecognized risk of nighttime caffeine use and provide new insights into the neural mechanism that mediates caffeine-induced changes in behavior.

## Results

### Nighttime Caffeine Consumption Impairs Inhibitory Control

To examine the effects of nighttime caffeine consumption on inhibitory control, we fed independent groups of *Canton-S* (*CS*) female and male flies food containing different caffeine concentrations (1, 5, 7.5, and 10 mg/ml)^20–22^. We then assessed their performance in the Go/No-Go test, which measures the ability to suppress movement in response to adverse conditions such as strong airflow or predator sounds, where movement could be detrimental to survival^18,23^. Typically, flies halt their movement in response to strong airflow, maintaining suppression until the airflow ceases. Flies with impaired inhibitory control exhibit impulsive flying, characterized by rapid movements at speeds exceeding 60 mm/sec. This behavior, quantified as a loss of inhibition event (LIE), serves as a measure of impulsivity^18,23^. In the absence of caffeine, both female and male flies showed strong movement suppression with negligible LIEs (Figure 1A-B, 0 mg/ml). However, caffeine-fed flies displayed a dose-dependent increase in LIEs with females exhibiting a more pronounced response than males (Figure 1A: Mann-Whitney; **, *p* < 0.005; ***, *p* < 0. 0.0001; *n* = 30-32. Figure 1B: Mann-Whitney; **, *p* < 0.005; ***, *p* < 0. 0.0001; *n* = 30-32).

**Figure 1.**
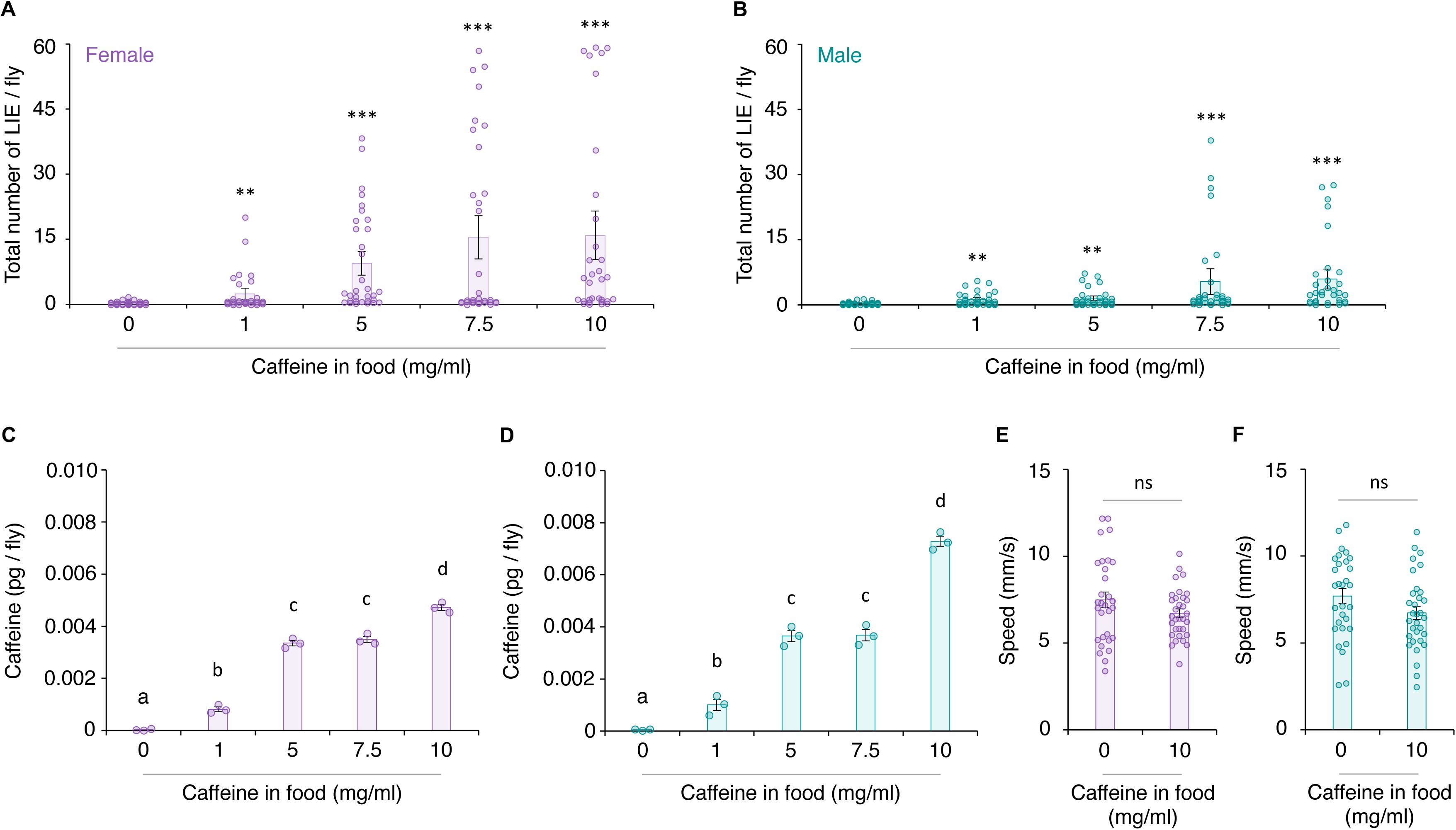
Nighttime caffeine consumption induces motor impulsivity in a dose-dependent manner. A-B. Nighttime caffeine intake increases motor impulsivity in wild-type *CS* females (A, purple bars) and males (B, teal bars) in a dose-dependent manner (Mann-Whitney; **, *p* < 0.005; ***, *p* < 0. 0.0001; *n* = 30-32). C-D. Caffeine levels per fly increase with higher caffeine concentrations in the food for both sexes. ANOVA with *post hoc* Tukey multiple comparison test; different letters denote statistically significant differences (*p* < 0.05; n = 3). E-F. Caffeine intake did not significantly affect baseline locomotor activity in females or males (Student *t*-test; ns, *p* > 0.05 *n* = 30-32).

To examine whether impulsivity correlates with caffeine levels in flies, *CS* flies were housed overnight on food containing varying concentrations of caffeine, and then caffeine levels were measured at the time of behavioral testing. As the caffeine dose increased, caffeine levels in both females and males rose accordingly (Figure 1C, female: ANOVA, *F_4,14_* = 443.97, *p* < 0.0001, *n* = 3; Figure 1D, male: ANOVA, *F_4,14_* = 229.09; *p* < 0.0001, *n* = 3), linking behavioral changes to internal caffeine levels. At 1, 5, or 7.5 mg/mL caffeine, females and males showed comparable caffeine content, but at 10 mg/mL caffeine, females had significantly lower caffeine levels than males (Figure 1C-D). Given that caffeine-induced impulsivity was greater in females than males (Figure 1A-B), these findings suggest that females are more sensitive to caffeine’s effects on impulsivity.

### Caffeine-Induced Impulsivity is Independent of Hyperactivity

To determine whether caffeine-induced impulsivity was associated with hyperactivity, we measured the walking speeds of *CS* females and males under baseline conditions (without airflow). Caffeine intake did not significantly affect walking speed in either females (Figure 1E: Student *t*-test, *p* = 0.149; *n* = 30–32) or males (Figure 1D: Student *t*-test, *p* = 0.070; n = 30–32). These findings indicate that the observed increase in impulsivity is not attributable to a general increase in locomotor activity.

### Sleep Deficits Do Not Account for Caffeine-Induced Impulsivity

Caffeine is known to promote wakefulness and disrupt sleep^24–26^. Consistent with this, we observed significant sleep reductions in caffeine-fed flies, with females showing greater sleep loss than males (Figure 2C: Student *t*-test: *, *p* < 0.05; ***, *p* < 0.0001; *n* = 14-15). To determine whether sleep loss contributes to motor impulsivity, we subjected *CS* flies to mechanical sleep disruption (MSD^27^) or light-induced sleep disruption (LSD^28^) during the nighttime phase and assessed their performance in the Go/No-Go assay. Neither MSD nor LSD increased LIEs in females or males (Figure 2A, female: Mann-Whitney, ns, *p* > 0.05; *n* = 14–17; Figure 2B: Mann-Whitney: ns, *p* > 0.05; *n* = 14–17). Although LSD led to significant sleep loss in females comparable to that seen with caffeine feeding, it had no effect in males (Figure 2D: Females: Student *t*-test: ***, *p* < 0.0001; *n* = 30; Males: Mann-Whitney: ns, *p* > 0.05; *n* = 29). These results indicate that caffeine-induced impulsivity cannot be explained solely by sleep deficits.

**Figure 2.**
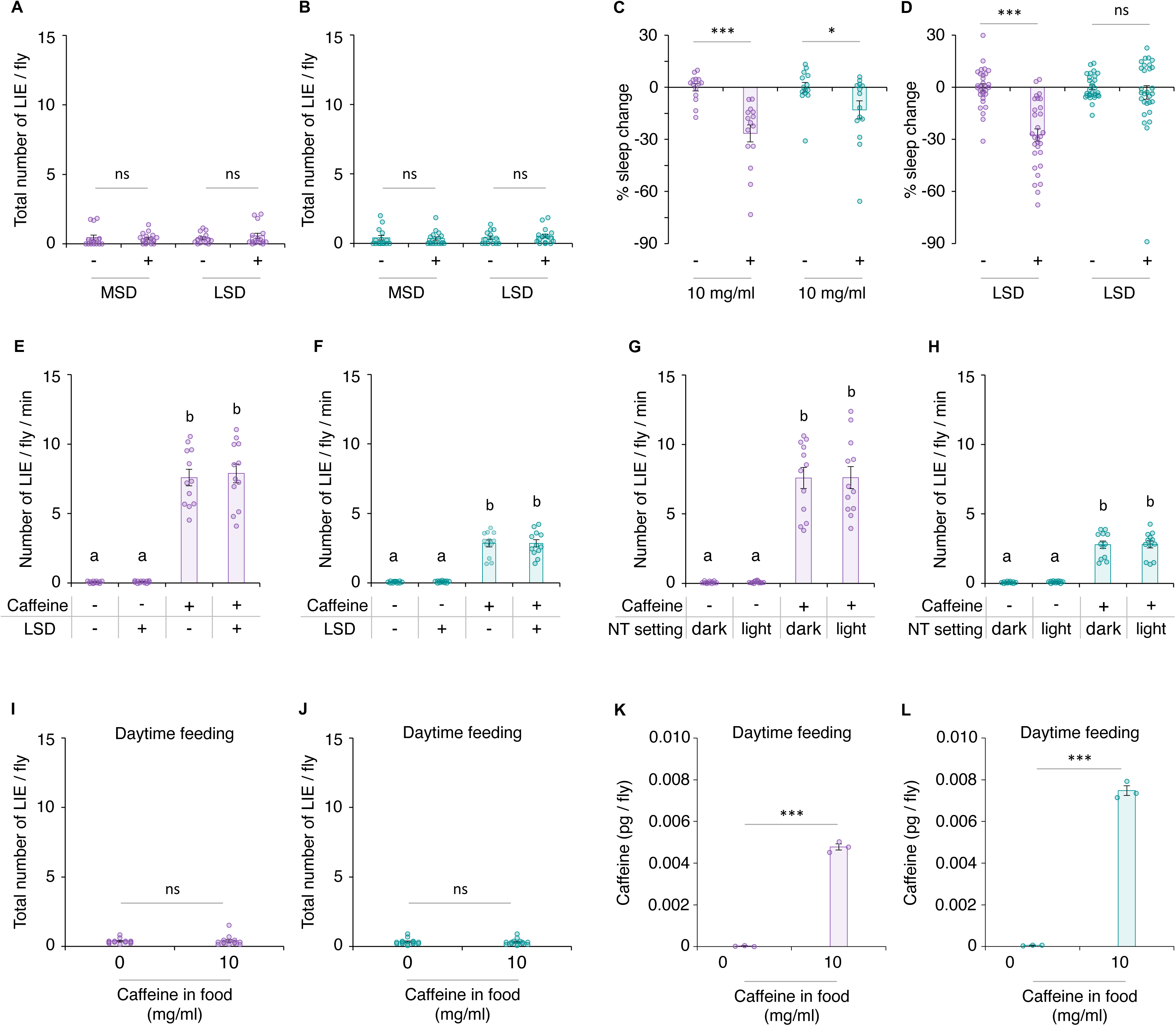
Nighttime sleep disruption or daytime caffeine feeding does not induce motor impulsivity. A-B. Neither nighttime mechanical sleep disruption (MSD; left panels in A and B) nor light-induced sleep disruption (LSD; right panels in A and B;) led to motor impulsivity in females or males (Mann-Whitney: ns, *p* > 0.05; *n* = 13-17). C. Caffeine-fed females and males showed comparable levels of sleep loss (Student *t*-test: *, *p* < 0.05; ***, *p* < 0.0001; *n* = 14-15). D. LSD induced significant sleep loss in females but not in males (For females, Student *t*-test; ***, *p* < 0.0001; *n* = 30; For males, Mann-Whitney; ns, *p* > 0.05; *n* = 29). E-F. LSD did not enhance caffeine-induced motor impulsivity in either sex (ANOVA with *post hoc* Tukey; *p* < 0.05; *n* = 12). G-H. Nighttime (NT) lighting conditions, either darkness or constant light, did not affect caffeine-induced motor impulsivity in females or males (ANOVA with *post hoc* Tukey; *p* < 0.05; *n* = 12). I-J. Daytime caffeine feeding for 4 h did not induce motor impulsivity in either sex (Mann-Whitney: ns, *p* > 0.05; *n* = 13-14). K-L. caffeine content per fly increased after 4 h of daytime feeding in both females and males (Student *t*-test or Mann-Whitney; ***, *p* < 0.0001; *n* = 3).

To further test this, we examined whether combining LSD and caffeine at night would exacerbate motor impulsivity. In both sexes, LSD did not enhance the impulsivity induced by nighttime caffeine intake (Figure 2E, female: ANOVA, *F_3,47_* = 96.88, *p* < 0.0001, *n* = 12; Figure 2F, male: ANOVA, *F_3,47_* = 76.80, *p* < 0.0001, *n* = 12), reinforcing the notion that caffeine’s effects are distinct from sleep loss. We also assessed whether nighttime lighting conditions influence caffeine-induced impulsivity. Flies exposed to constant darkness or light during caffeine intake exhibited comparable impulsivity levels (Figure 2G, female: ANOVA, *F_3,47_* = 62.31, *p* < 0.0001, *n* = 12; Figure 2H, male: ANOVA, *F_3,47_* = 68.94, *p* < 0.0001, *n* = 12). Together, these findings suggest that caffeine’s effects on motor impulsivity are independent of both sleep loss and environmental light conditions at night.

### Daytime Caffeine Intake Does Not Impair Inhibitory Control

To determine whether caffeine’s effect on impulsivity depends on the timing of intake, we fed flies with 10 mg/ml caffeine for 4 h during the daytime and assessed their inhibitory control. The 4-h daytime feeding regime was selected because it resulted in caffeine levels (Figure 2K-L) comparable to those observed in flies fed caffeine at night (Figure 1C-1D). Both females and males fed with caffeine during the daytime exhibited robust inhibitory control (Figure 2I-J, female: Mann-Whitney, *p* > 0.05, *n* = 13-14; Figure 2J, male: Mann-Whitney, *p* > 0.05, *n* = 14). These results clearly indicate that it is the timing, specifically nighttime, of caffeine intake that leads to impaired inhibitory control and motor impulsivity.

### Dopamine Mediates Caffeine-Induced Impulsivity

Caffeine’s wake-promoting effects are known to involve dopaminergic signaling^24,26^. To determine whether dopamine also mediates caffeine’s impact on inhibitory control, we examined flies carrying heterozygous mutations in *pale* (*ple/+*; two independent alleles), which encodes tyrosine hydroxylase, the rate-limiting enzyme in dopamine biosynthesis^29^. In the absence of caffeine, *ple/+* displayed pronounced movement inhibition (data not shown). Upon caffeine feeding, *ple/+* mutants retained strong movement suppression, unlike control *CS* flies (Figure 3A, female: ANOVA, *p* = 0.001, *n* = 5-6; Figure 3B, male: ANOVA, *p* < 0.0001, *n* = 6), suggesting that reduced dopamine biosynthesis blocks caffeine-induced impulsivity.

**Figure 3.**
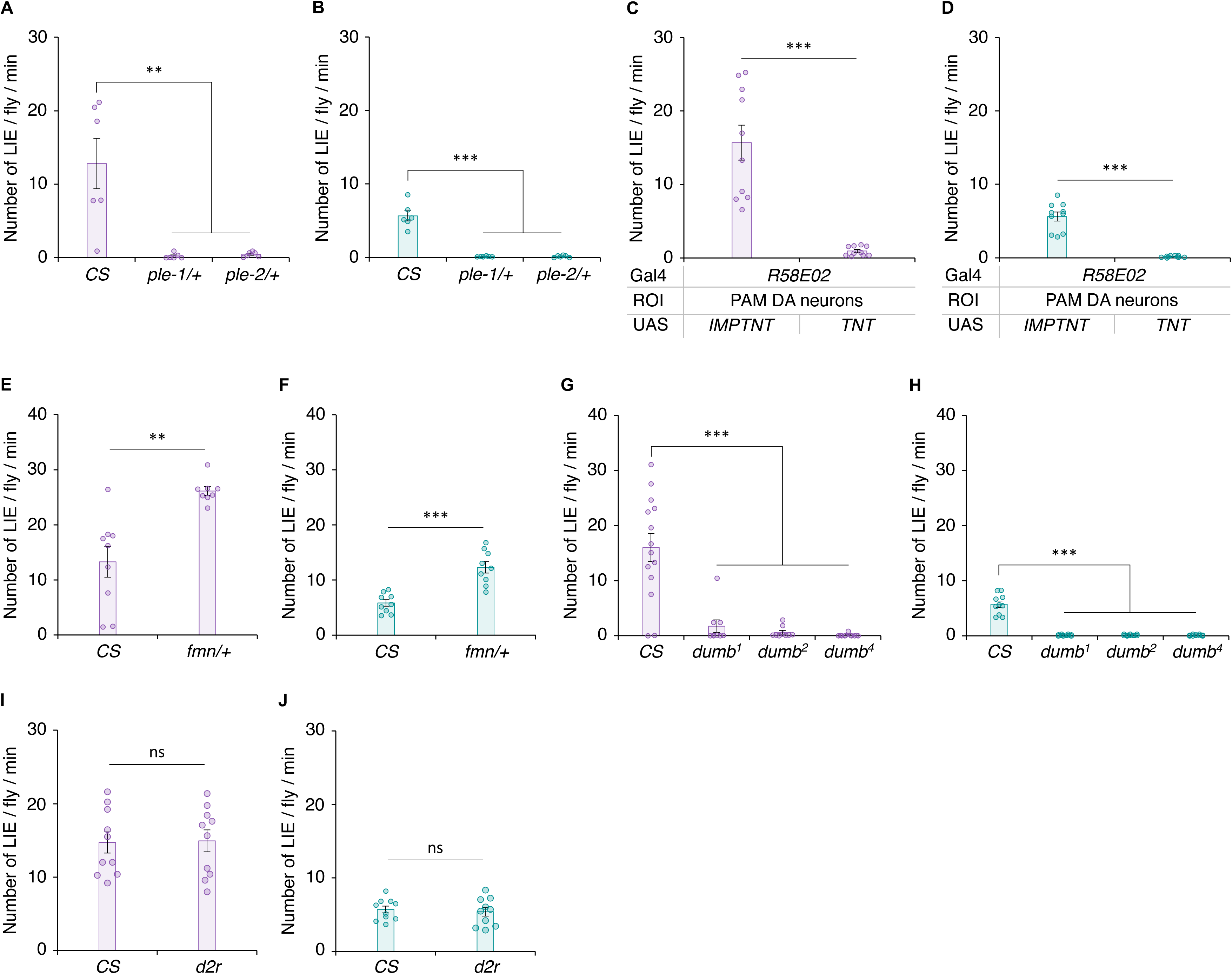
Caffeine-induced impulsivity requires PAM dopaminergic neurons and D1 receptor signaling. A-B. Caffeine-fed *pale* (*ple*) heterozygotes (*ple-1/+* and *ple-2/+;* purple bars – female; teal bars – male) exhibited robust movement inhibition (ANOVA with *post hoc* Dunnett; **, *p* < 0.005, ***, *p* < 0.0001; *n* = 5-6). C-D. Silencing PAM dopaminergic neurons using via tetanus toxin (TNT) light chain (*R58E02-GAL4/UAS-TNT*) abolished caffeine-induced impulsivity (Student *t*-test; ***, *p* < 0.0001; *n* = 10). E-F. *fumin* (*fmn*) heterozygotes (*fmn/+*) displayed significantly heightened impulsivity after caffeine feeding (Student *t*-test; **, *p* < 0.005; ***, *p* < 0.0001; *n* = 8-9). G-H. Three D1 (dDA1) receptor mutant alleles (*dumb^1^*, *dumb^2^* and *dumb^4^*) showed no increase in impulsivity following caffeine feeding (ANOVA with post hoc Dunnett using *CS* as a control: ***, *p* < 0.0001; *n* = 9-14). I-J. The D2 receptor mutant (*d2R*) showed a caffeine response similar to *CS* (Student *t*-test: ns, *p* > 0.05; *n* = 10).

To further assess the role of dopamine, we silenced PAM dopaminergic neurons, previously linked to caffeine-induced arousal^24^, using tetanus toxin (TNT). Silencing PAM neurons similarly abolished caffeine-induced impulsivity (Figure 3C, female: Student’s *t*-test, ***, *p* < 0.0001, n = 10; Figure 3D, male: Student’s *t*-test, ***, *p* < 0.0001, n = 10), supporting the involvement of this neural cluster.

As a complementary approach, we tested flies heterozygous for the dopamine transporter mutation *fumin* (*fmn/+*), which increases extracellular dopamine^30^. Upon caffeine feeding, *fmn/+* flies exhibited significantly elevated impulsivity levels (Figure 3E, female: Student *t*-test, *p* = 0.002, *n* = 8-9; Figure 3F, male: Student *t*-test, *p* < 0.0001, *n* = 9), further implicating dopamine in mediating caffeine’s effect on inhibitory control.

**Figure 4.**
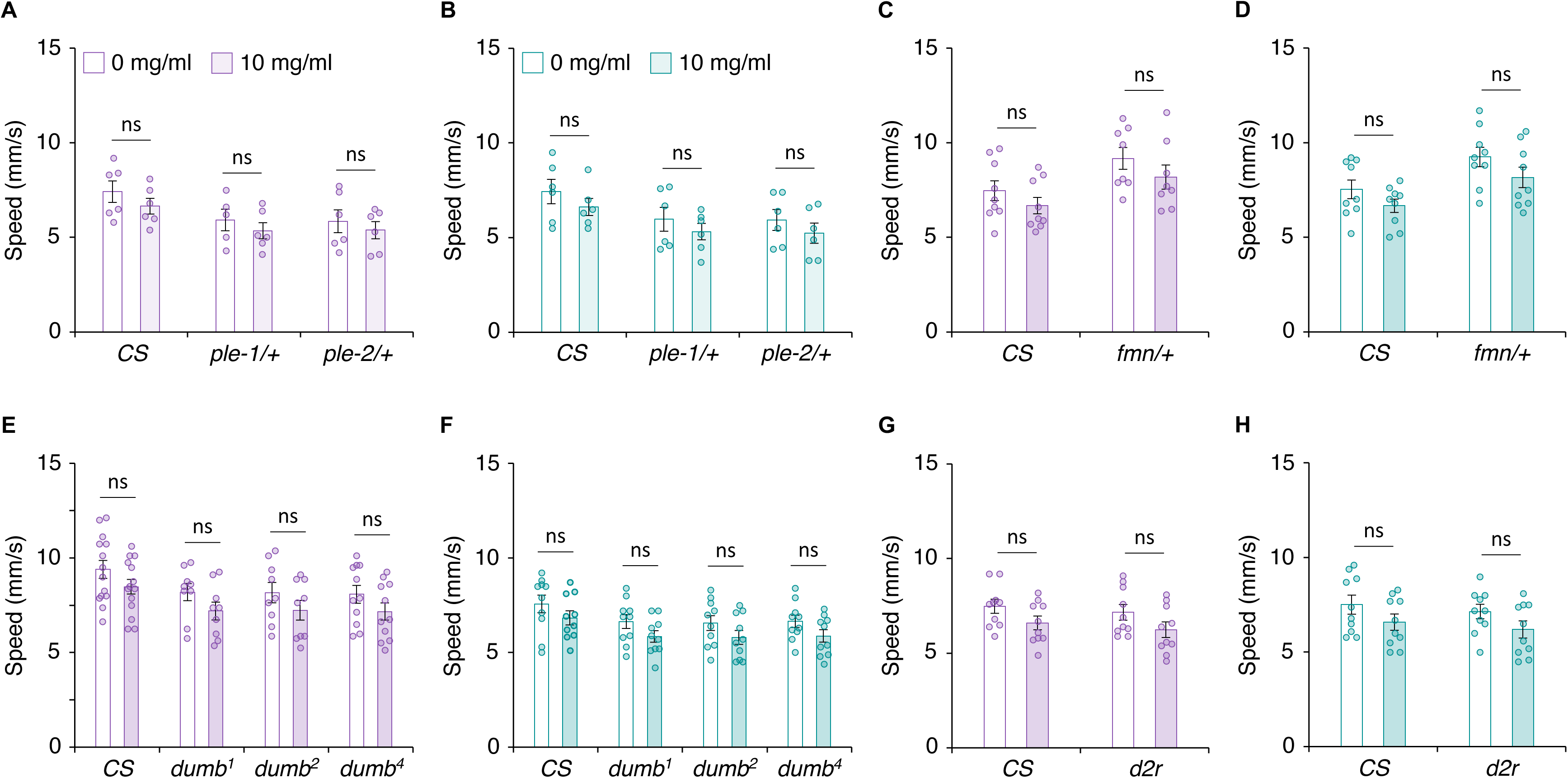
Nighttime caffeine feeding does not alter basal locomotor activity at the time of GNG testing. Caffeine-fed *ple/+* (A-B; *n* = 5-6), *fmn/+* (C-D; *n* = 8-9), D1 receptor mutants *dumb^1^*, *dumb^2^* and *dumb^4^* (E-F; *n* = 9-14) and D2 receptor mutant *d2r* (G-H; *n* = 10) exhibited basal locomotor activities comparable to unfed controls (Student *t*-test: ns, *p* > 0.05; *n* = 5-14).

### Dopamine D1 Receptor Mediates Caffeine-Induced Impulsivity

The dopamine D1 receptor, dDA1, also known as Dop1R1, has previously been identified as a key mediator of caffeine’s wake-promoting effects^26^. To determine its role in caffeine-induced impulsivity, we examined three independent *dDA1* mutant alleles (*dumb^1^*, *dumb^2^*, and *dumb^4^*)^19,31^. None of these mutants exhibited impulsivity following caffeine feeding, in contrast to wild-type controls (Figure 3G, female: ANOVA, *p* < 0.0001, *n* = 9-14; Figure 3H, male: ANOVA, *p* < 0.0001, *n* = 10). In comparison, D2 receptor mutants (*d2r*)^31^ responded similarly to wild-type *CS*, showing impulsivity after caffeine feeding (Figure 3I, female: Student’s *t*-test, ns, *p* > 0.05; *n* = 10; Figure 3I, male: Student’s *t*-test, ns, *p* > 0.05; *n* = 10). These findings indicate that D1/dDA1, but not D2/d2R, mediates caffeine-induced impulsivity. Importantly, caffeine-induced impulsivity was not associated with hyperactivity. Walking speeds of all dopamine mutants – *ple/+* (4A-B; Student *t*-test: ns, *p* > 0.05; *n* = 5-6), *fmn/+* (4C-D; Student *t*-test: ns, *p* > 0.05; *n* = 8-9), *dDA1* mutant alleles *dumb^1^*, *dumb^2^* and *dumb^4^* (4E-F; Student *t*-test: ns, *p* > 0.05; *n* = 9-14) and D2 mutant *d2r* (4G-H; Student *t*-test: ns, *p* > 0.05; *n* = 10) – remained comparable to their unfed controls, like *CS*.

dDA1 is highly expressed in mushroom body (MB) neurons, which regulate complex behaviors such as learning, and memory^32^. The MB consists of three primary substructures: γ-lobes, α/β-lobes, and α’/β’-lobes. To dissect the contributions of these substructures to impulsivity, we knocked down dDA1 in each lobe using split-GAL4 drivers: MB005B (α’/β’), MB008B (α/β) and MB009B (γ). Knockdown of dDA1 in γ or α/β lobes significantly suppressed caffeine-induced impulsivity, whereas knockdown in α’/β’ lobes had no effect (Figure 5A, female: ANOVA with *post hoc* Dunnett, *p* < 0.0001, *n* = 10; Figure 5B, male: ANOVA with *post hoc* Dunnett, *p* < 0.0001, *n* = 10).

**Figure 5.**
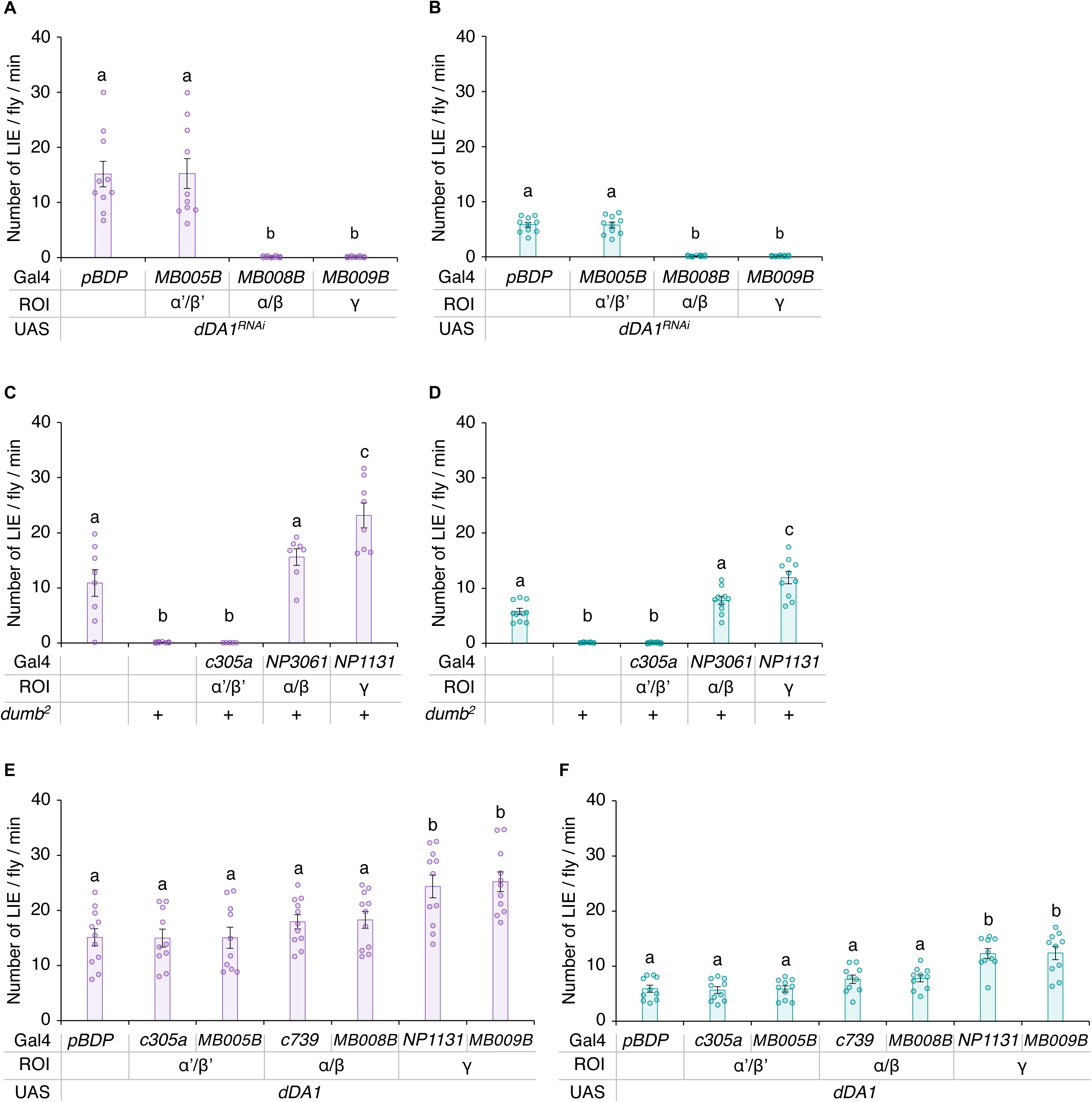
Dopamine D1 receptor (dDA1) in the mushroom body α/β and γ neurons mediates caffeine-induced impulsivity. A-B. Knockdown of *dDA1* (*UAS-dDA1 RNAi*) in α/β (*MB008B*) or γ (*MB009B*) neurons, but not in α’/β’ (MB005B) neurons, abolished caffeine-induced impulsivity (ANOVA with *post hoc* Dunnett, *p* < 0.0001; *n* = 10; different letters indicate statistically significant differences: purple bars, females; teal bars, males). C-D. Reinstatement of dDA1 in α/β (*NP3061,dumb^2^*) or γ (*NP1131;dumb^2^*) neurons in *dumb^2^* restored caffeine-induced impulsivity while reinstatement in α’/β’ (*c305a;dumb^2^*) neurons did not (ANOVA with *post hoc* Tukey; ns, *p* > 0.05, **, *p* < 0.005, ***, *p* < 0.0001; *n* = 6-10). E-F. Overexpression of dDA1 in α’/β’ (*c305a* and *MB005B*) or α/β (*c739* and *MB008B*) neurons resulted in caffeine-induced impulsivity comparable to control (*pBDP*), whereas overexpression in γ (*NP1131* and *MB009B*) neurons led to significantly higher impulsivity (ANOVA with *post hoc* Dunnett, ***, *p* < 0.0001; *n* = 10-11).

To further test the lob-specific requirement of dDA1, we reinstated its expression in individual MB lobes in the *dumb^2^* mutant background^19,33^. Restoring dDA1 in γ lobes (NP1131) or α/β lobes (NP3061) fully rescued caffeine-induced impulsivity, whereas restoration in α’/β’ lobes (c305a) did not (Figure 5C, female: ANOVA, *p* < 0.0001, *n* = 6–8; Figure 5D, male: ANOVA, *p* < 0.0001, *n* = 10). Interestingly, γ-lobe restoration led to even greater impulsivity than that observed in *CS* or in α/β-lobe rescue, suggesting heightened sensitivity of γ-lobe neurons to dDA1 activity. To test this directly, we overexpressed dDA1 in individual MB lobes in a wild-type background. Overexpression in γ lobes (NP1131 or MB009B) significantly enhanced caffeine-induced impulsivity relative to *CS,* whereas overexpression in α/β lobes (c739 or MB008B) or α’/β’ lobes (via c305a or MB005B) had no effect (Figure 5E, female: ANOVA, *p* < 0.0001, *n* = 10–11; Figure 5F, male: ANOVA, *p* < 0.0001, *n* = 10). Together, these findings demonstrate that dDA1 in the α/β and γ lobes of the MB is essential for mediating caffeine-induced impulsivity, with the γ lobes exhibiting heightened sensitivity to dDA1 activity.

## Discussion

In this report, we have shown that nighttime caffeine intake disrupts inhibitory control in *Drosophila* through dopamine D1 receptor (dDA1) signaling in the MB, revealing a neural and molecular framework by which caffeine modulates impulse control. Notably, this behavioral phenotype was not attributable to general hyperactivity, reinforcing its specificity to inhibitory control. These findings expand the behavioral repertoire influenced by caffeine beyond arousal and provide a mechanistic basis for its time-dependent effects on cognitive function.

We show that caffeine impairs inhibitory control only when consumed at night, despite similar internal caffeine concentrations during day and night intake. This finding suggests a strong interaction between caffeine’s effects and temporal factors, likely involving circadian regulation. On the contrary, sleep deprivation alone, whether mechanical or light-induced, did not produce impulsivity, nor did it amplify caffeine’s effects. These results together suggest that caffeine-induced impulsivity is independent of generalized arousal or sleep loss, implicating a distinct, temporally gated mechanism. This chrono-dependent sensitivity parallels findings from human studies, where cognitive and behavioral responses to caffeine vary by circadian phase. For example, morning, but not evening, caffeine intake promotes performance enhancements, while evening caffeine intake delays melatonin rhythms and disrupts sleep architecture^20–22^. These findings underscore the significance of circadian context in shaping caffeine’s impact on behavior.

Our work establishes that dopaminergic signaling is necessary and sufficient for caffeine’s effect on inhibitory control. Caffeine-induced impulsivity was abolished in flies with reduced dopamine biosynthesis or silenced PAM dopaminergic neurons, and conversely, it was amplified in flies with elevated extracellular dopamine. Targeted manipulations in MB further identified the dDA1 receptor in the α/β and γ lobes as critical for mediating this dopaminergic effect. Consistent with our findings, D1 receptor signaling in the prefrontal cortex is known to regulate impulse control in both rodents and humans, and caffeine has been shown to impair decision-making and promote risk-taking behavior^34–39^. Notably, high caffeine consumption in children, adolescents and young adults has been linked to greater sensation seeking and risk-taking tendencies^40,41^. In gamblers, caffeine use is associated with earlier onset of gambling, heightened impulsivity, and increased nicotine co-use^37^. Also in spontaneously hypertensive rats, an animal model of ADHD, chronic caffeine treatment corrects dopaminergic dysfunction and improves attention and working memory, supporting a functional link between caffeine and dopamine signaling in cognitive control^42^.

Caffeine promotes wakefulness in rodents by antagonizing adenosine receptors and enhancing dopamine signaling^43,44^. In *Drosophila*, it enhances wakefulness independently of adenosine antagonism^25^ but requires dopamine signaling. Specifically, caffeine acts presynaptically to enhance dopaminergic activity in PAM neurons^24^. Its wake-promoting effect is then mediated postsynaptically by dDA1 in the MB α/β and γ lobes^26^. Notably, our results suggest that the same neural circuitry also mediates caffeine-induced motor impulsivity independently of sleep loss. This MB substructures are also known to regulate multiple complex behaviors such as learning, memory, behavioral flexibility, and decision making^16–19,23,45–48^. Future research should explore how caffeine’s diverse effects are mediated by overlapping neurochemical pathways and neural structures.

Our findings align with growing evidence that dopaminergic signaling is closely intertwined with circadian timing. Dopamine release follows daily rhythms, peaking during the active phase in both flies and rodents, and is itself regulated by, and contributes to, circadian clock function^49,50^. In *Drosophila*, light-dependent upregulation of dDA1 in specific circadian neurons promotes morning arousal, highlighting time-of-day-dependent dopamine sensitivity^50^. This relationship is reciprocal as dopamine modulates circadian gene expression and neuronal excitability, and its disruption contributes to disorders like Parkinson’s disease, which features both circadian and dopaminergic dysfunction^51,52^. Notably, PAM cluster dopamine neurons implicated in caffeine-induced impulsivity also show circadian variation in vulnerability to oxidative stress, which is abolished in clock gene mutants^53^. Together, these data suggest that caffeine acts on a circadian-gated dopaminergic system that regulates behavioral inhibition. Temporal sensitivity of dopamine signaling may underlie the time-specific effects of caffeine on impulsivity.

Another novel aspect of our findings is the identification of sex differences in caffeine-induced impulsivity, with female flies displaying greater sensitivity than males. This mirrors human studies showing that females are more likely to experience negative caffeine-related effects such as anxiety and nervousness, whereas males more often report positive effects, like enhanced perception and vigor^54,55^. Similar sex-specific responses to caffeine have been seen in rodents^56–60^; for example, regular intake of caffeinated coffee induces sex-dependent changes in mice, with females exhibiting increased dominance and self-care and male showing greater sociability^60^. Together, these findings underscore the complex, sex-dependent nature of caffeine’s behavioral effects. Our study provides a tractable model for uncovering the neural and molecular mechanisms underlying sex-specific sensitivity to caffeine, particularly in the context of impulsivity.

In summary, our study shows that nighttime caffeine intake disrupts inhibitory control via D1 (dDA1) dopamine receptor signaling in the MB α/β and γ lobes, with effects that are independent of sleep deprivation and modulated by both sex and circadian phase. These findings advance our understanding of how executive function is influenced by caffeine and underscore *Drosophila*’s utility for dissecting the neural basis of impulsivity. Importantly, we provide direct evidence that the behavioral impact of psychoactive substances, like caffeine, is time-of-day dependent, providing support to chronotherapy approaches and emphasizing the need to integrate circadian biology into behavioral neuroscience.

## Acknowledgments

We are grateful to the Bloomington Stock Center and Drs. Dubnau and Anderson for sharing fly lines, the Cytometry, Screening, and Imaging Core at Border Biomedical Research Center on confocal microscopy. We are also thankful to the past and current lab members for their help, discussion, and support. This work was supported by the National Institutes of Health grant R21MH109953 (KAH), Brain & Behavior Research Foundation grant (KAH) and National Institutes of Health grant R16GM145548(KAH).

National Institutes of Health grant R21MH109953 (KAH)

Brain & Behavior Research Foundation grant (KAH)

National Institutes of Health grant R16GM145548(KAH)

## Author contributions

Conceptualization: EBS, PRS, KAH

Methodology: EBS, PRS

Investigation: EBS, PRS

Visualization: EBS, PRS

Supervision: KAH

Writing—original draft: EBS, KAH

Writing—review & editing: EBS, PRS, KAH

## Data and materials availability

All raw data and materials used in the study are available upon request from the corresponding authors.

## Footnote

Erick B. Saldes current address: Department of Cancer Biology & Pharmacology, The University of Illinois College of Medicine Peoria, One Illini Dr. Peoria, IL 61615

## STAR METHODS

KEY RESOURCES TABLE

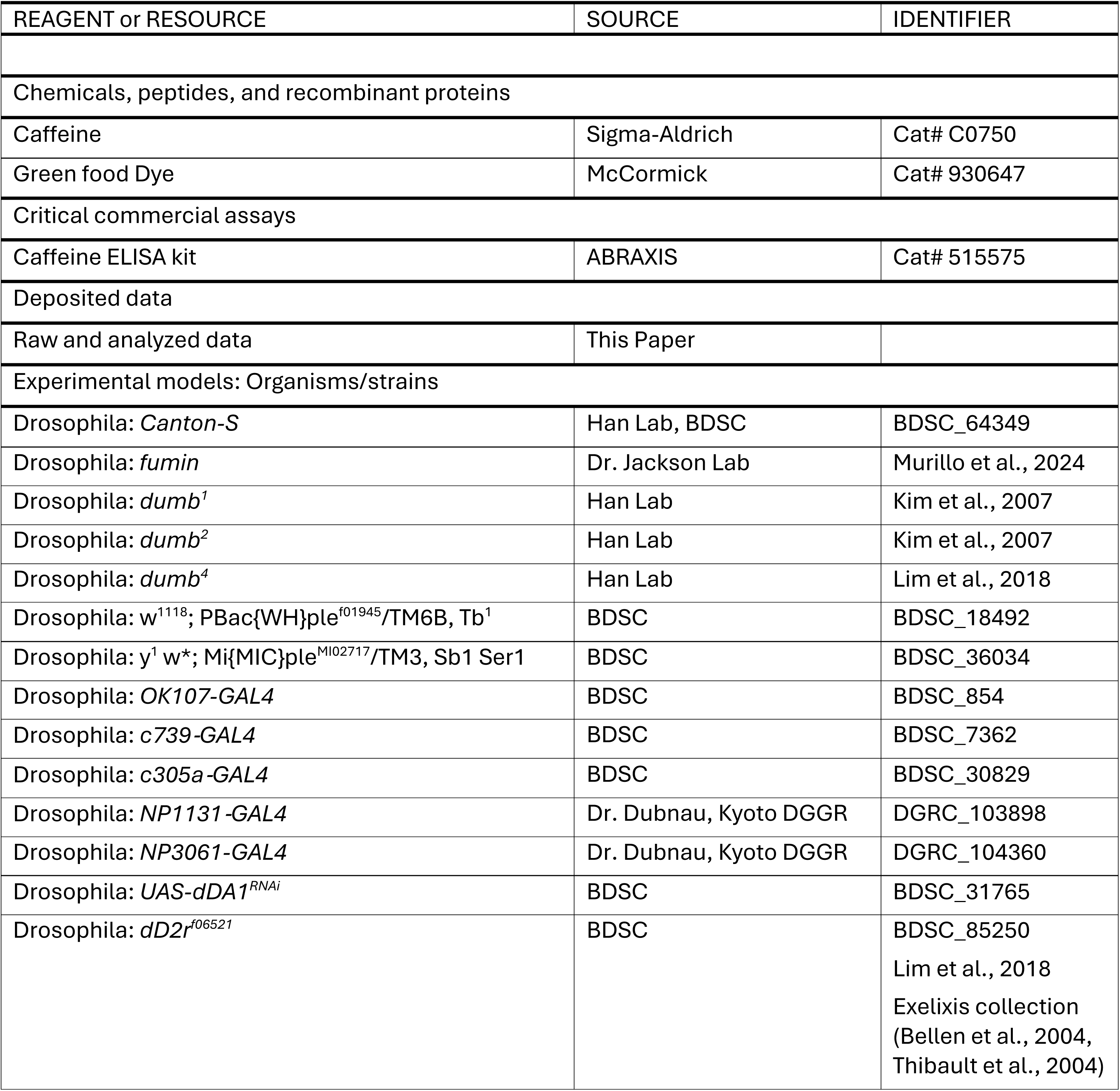
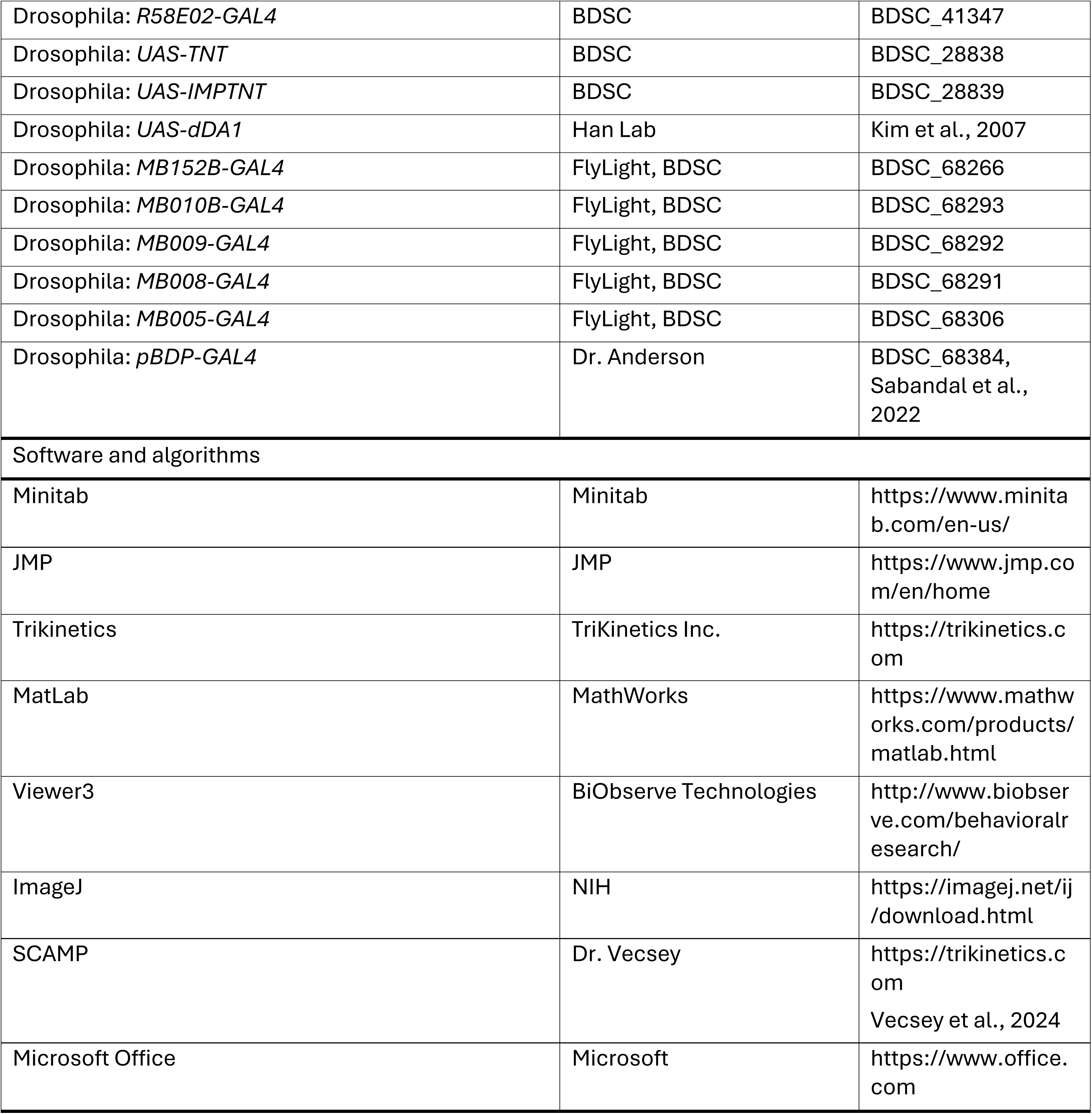

### RESOURCE AVAILABILITY

#### Lead contact

Any information, resource, and reagent requests should be directed to the lead contact, Kyung-An Han.

#### Materials availability

This study did not generate new reagents and most fly strains used in the study are available from the Bloomington Drosophila Stock Center. The D1 receptor mutant alleles and *UAS-dDA1* fly lines are available on request from the lead contact.

#### Data and code availability

All data reported in this study is available from the lead contact upon request.

### EXPERIMENTAL MODEL AND STUDY PARTICIPANT DETAILS

In this study, we used *Drosophila melanogaster* as a model system. Specific information about fly strains, culture, and maintenance as well as specific experimental conditions are found in the subsequent sections below.

### METHOD DETAILS

#### Fly strains and culture

The wild-type strain used in this study is *Canton-S* (*CS*). We previously described *fumin* (*fmn*)^61^, *dumb^1^* ^19^, *dumb^2^* ^19^, *dumb^4^* ^31^, *d2r*^31^, and *UAS-dDA1*^19^ – all of which were placed in the *CS* background. The following stocks were obtained from the Bloomington Drosophila stock center (Bloomington, IN): *ple^f01945^*(noted as ple-1; 18492), *ple^MI02717^* (noted as ple-2; 36034), *UAS-TNT* (28838), *UAS-IMP-TNT* (28839), *UAS-dDA1 RNAi* (31765), *GMR58E02-GAL4* (41347), *c305a-GAL4* (30829), *MB005B-GAL4* (68306), *MB008B-GAL4* (68291), and *MB009B-GAL4* (68292); *pBDP-GAL4* from Dr Anderson (California Institute of Technology, Pasadena, CA) and *NP1131-GAL4* and *NP3061-GAL4* from Dr. Dubnau (Stony Brook University School of Medicine, Stony Brook, NY). All *Drosophila* strains were raised on a standard cornmeal/sucrose/yeast/agar medium at 25° C with 50 % relative humidity under a 12 h light/12 h dark cycle. For experiments, female and male flies were collected under carbon dioxide within two days after eclosion and housed in same-sex groups of 13 per vial (representing n = 1). Both females and males were examined separately.

#### Caffeine feeding

Caffeine (Sigma-Aldrich, C0750) was mixed into melted standard fly food medium (cornmeal/sucrose/yeast/agar) at a concentration of 10 mg/mL for all experiments except for the dose-response analysis in Figure 1A and 1B, where additional concentrations of 1, 5 and 7.5 mg/mL were used^18,26,62–65^. For nighttime caffeine exposure, groups of 13 female or male flies (n = 1) were transferred to vials containing caffeine-laced food 30 min before lights-off and remained on this food throughout the dark cycle (ZT12–ZT24). Flies were immediately tested in the Go/No-Go assay at the end of the dark period. For the daytime feeding condition, flies were exposed to 10 mg/mL caffeine for 4 h (ZT0–ZT4) and tested immediately afterward. To confirm ingestion of the caffeine-containing food, green food dye (McCormick, 930647) was added to the mixture of fly food medium and caffeine, and consumption was verified by visual inspection of dye accumulation in the abdomen.

#### Sleep Disruption

Two distinct methods were used to disrupt sleep in *Drosophila*. For mechanical sleep deprivation (MSD), vials containing flies were placed on a vortexer and subjected to randomized shaking for 2 s within every 20-s interval throughout the dark cycle, as previously described^27^. For light-induced sleep disruption (LSD), lights were randomly turned on for 30 min once every hour during the dark phase, following established protocols^28^. Flies were immediately subjected to the Go/No-Go test at the end of the dark cycle to assess behavioral consequences of sleep disruption.

#### Go/No-Go test

The Go/No-Go test was performed as previously described^18,23^. Briefly, Flies were placed individually into a clear rectangular plexiglass chamber (60 mm L X 60 mm W X 15 mm H) connected to a filtered air source. After a 1-min acclimation period to explore the chamber, a continuous airflow of 10 L/min was applied for 10 min. Fly behavior was video recorded to monitor movements before and during airflow exposure. Videos were analyzed for flying events manually or using Viewer3 tracking software (BIObserve Technologies, Bonn, Germany), which enables automated tracking and quantification of individual fly movement speed in mm/sec. Raw data were exported to Excel (Microsoft, Redmond, WA). Movements exceeding 60 mm/sec were classified as loss of inhibition events (LIEs^18,23^), which were scored per fly per min. Both total LIE or peak LIEs (maximum LIEs per minute) were used for comparison between control and experimental groups. Baseline locomotor activity during the initial no-airflow phase (“go” period) was assessed using average walking speed (reported in Figure 1E-F and Figure 3A-H). All behavioral experiments were conducted with the experimenter blind to condition, sex, and genotype. Control and experimental groups were always tested within the same experimental session. Multiple independent groups of flies derived from separate seedings, or genetic crosses were used for all behavioral assays.

#### Caffeine analysis

Caffeine levels in flies were measured using a commercially available Caffeine ELISA kit (Catalog no. 515575; ABRAXIS, Warminster, PA, USA). For each feeding condition and sex, five whole flies were collected and transferred into an ice-cold KONTES Micro Tissue Grinder (ThermoFisher Scientific, Waltham, MA). Flies were homogenized in 50 μl of Sample Diluent flies for 30 – 45 s. The homogenate was then transferred to a clean microcentrifuge tube and centrifuged at 14,000 rpm for 10 min at 4° C. The resulting supernatant was used for caffeine quantification according to the manufacturer’s instructions. Three independent biological replicates (each from separate broods and feedings) were analyzed per treatment group and sex.

### QUANTIFICATION AND STATISTICAL ANALYSIS

All statistical analyses were performed using either the Minitab software (Minitab, State College, PA) or JMP (SAS, Cary, NC). Raw data were analyzed using the Anderson–Darling goodness-of-fit test for distribution and are reported as mean + SEM. Normally distributed data were analyzed by either a two-tailed Student’s *t*-test for two groups or by ANOVA followed by *post hoc* Tukey’s multiple-comparison for three or more groups or Dunnett’s test to compare experimental groups with a control group. Non-normally distributed data were analyzed by Kruskal-Wallis and post hoc Mann-Whitney tests. Significant difference among the groups under comparison was determined using an α level of 0.05 in all analyses.

## References

1. Fulgoni, V.L., Keast, D.R., and Lieberman, H.R. (2015). Trends in intake and sources of caffeine in the diets of US adults: 2001-2010. Am J Clin Nutr 101, 1081–1087. 10.3945/ajcn.113.080077.

2. Walsh, J.K., Muehlbach, M.J., Humm, T.M., Dickins, Q.S., Sugerman, J.L., and Schweitzer, P.K. (1990). Effect of caffeine on physiological sleep tendency and ability to sustain wakefulness at night. Psychopharmacology (Berl) 101, 271–273. 10.1007/BF02244139.

3. Muehlbach, M.J., and Walsh, J.K. (1995). The effects of caffeine on simulated night-shift work and subsequent daytime sleep. Sleep 18, 22–29. 10.1093/sleep/18.1.22.

4. Lieberman, H.R., Tharion, W.J., Shukitt-Hale, B., Speckman, K.L., and Tulley, R. (2002). Effects of caffeine, sleep loss, and stress on cognitive performance and mood during US Navy SEAL training. Psychopharmacology 164, 250–261.

5. Franke, A.G., Bagusat, C., McFarlane, C., Tassone-Steiger, T., Kneist, W., and Lieb, K. (2015). The Use of Caffeinated Substances by Surgeons for Cognitive Enhancement. Ann Surg 261, 1091–1095. 10.1097/SLA.0000000000000830.

6. Ogeil, R.P., Barger, L.K., Lockley, S.W., O’Brien, C.S., Sullivan, J.P., Qadri, S., Lubman, D.I., Czeisler, C.A., and Rajaratnam, S.M.W. (2018). Cross-sectional analysis of sleep-promoting and wake-promoting drug use on health, fatigue-related error, and near-crashes in police officers. BMJ Open 8, e022041. 10.1136/bmjopen-2018-022041.

7. Irwin, C., Khalesi, S., Desbrow, B., and McCartney, D. (2020). Effects of acute caffeine consumption following sleep loss on cognitive, physical, occupational and driving performance: A systematic review and meta-analysis. Neurosci Biobehav Rev 108, 877–888. 10.1016/j.neubiorev.2019.12.008.

8. Bonnet, M.H., and Arand, D.L. (1994). Impact of naps and caffeine on extended nocturnal performance. Physiol Behav 56, 103–109. 10.1016/0031-9384(94)90266-6.

9. Kamimori, G.H., McLellan, T.M., Tate, C.M., Voss, D.M., Niro, P., and Lieberman, H.R. (2015). Caffeine improves reaction time, vigilance and logical reasoning during extended periods with restricted opportunities for sleep. Psychopharmacology (Berl) 232, 2031–2042. 10.1007/s00213-014-3834-5.

10. Wingelaar-Jagt, Y.Q., Bottenheft, C., Riedel, W.J., and Ramaekers, J.G. (2023). Effects of modafinil and caffeine on night-time vigilance of air force crewmembers: A randomized controlled trial. J Psychopharmacol 37, 172–180. 10.1177/02698811221142568.

11. Killgore, W.D., Kamimori, G.H., and Balkin, T.J. (2014). Caffeine improves the efficiency of planning and sequencing abilities during sleep deprivation. J Clin Psychopharmacol 34, 660–662. 10.1097/JCP.0000000000000184.

12. Huffmyer, J.L., Kleiman, A.M., Moncrief, M., Scalzo, D.C., Cox, D.J., and Nemergut, E.C. (2020). Impact of Caffeine Ingestion on the Driving Performance of Anesthesiology Residents After 6 Consecutive Overnight Work Shifts. Anesth Analg 130, 66–75. 10.1213/ANE.0000000000004252.

13. Daly, J.W., and Fredholm, B.B. (1998). Caffeine—an atypical drug of dependence. Drug and alcohol dependence 51, 199–206.

14. Parry, D., Iqbal, S., Harrap, I., Oeppen, R.S., and Brennan, P.A. (2023). Caffeine: benefits and drawbacks for technical performance. Br J Oral Maxillofac Surg 61, 198–201. 10.1016/j.bjoms.2023.01.007.

15. Jacobson, B.H., Winter-Roberts, K., and Gemmell, H.A. (1991). Influence of caffeine on selected manual manipulation skills. Percept Mot Skills 72, 1175–1181. 10.2466/pms.1991.72.3c.1175.

16. Cohn, R., Morantte, I., and Ruta, V. (2015). Coordinated and Compartmentalized Neuromodulation Shapes Sensory Processing in Drosophila. Cell 163, 1742–1755. 10.1016/j.cell.2015.11.019.

17. Modi, M.N., Rajagopalan, A.E., Rouault, H., Aso, Y., and Turner, G.C. (2023). Flexible specificity of memory in. Elife 12. 10.7554/eLife.80923.

18. Sabandal, P.R., Kim, Y.C., Sabandal, J.M., and Han, K.A. (2025). Social context and dopamine signaling converge in the mushroom body to drive impulsivity. bioRxiv. 10.1101/2025.02.21.639508.

19. Kim, Y.C., Lee, H.G., and Han, K.A. (2007). D1 dopamine receptor dDA1 is required in the mushroom body neurons for aversive and appetitive learning in Drosophila. J Neurosci 27, 7640–7647. 10.1523/JNEUROSCI.1167-07.2007.

20. Stojanović, E., Scanlan, A.T., Milanović, Z., Fox, J.L., Stanković, R., and Dalbo, V.J. (2022). Acute caffeine supplementation improves jumping, sprinting, and change-of-direction performance in basketball players when ingested in the morning but not evening. Eur J Sport Sci 22, 360–370. 10.1080/17461391.2021.1874059.

21. Zhang, Y., Yang, W., Xue, Y., Hou, D., Chen, S., Xu, Z., Peng, S., Zhao, H., Wang, C., and Liu, C. (2024). Timing Matters: Time of Day Impacts the Ergogenic Effects of Caffeine-A Narrative Review. Nutrients 16. 10.3390/nu16101421.

22. Burke, T.M., Markwald, R.R., McHill, A.W., Chinoy, E.D., Snider, J.A., Bessman, S.C., Jung, C.M., O’Neill, J.S., and Wright, K.P. (2015). Effects of caffeine on the human circadian clock in vivo and in vitro. Sci Transl Med 7, 305ra146. 10.1126/scitranslmed.aac5125.

23. Sabandal, P.R., Saldes, E.B., and Han, K.A. (2022). Acetylcholine deficit causes dysfunctional inhibitory control in an aging-dependent manner. Sci Rep 12, 20903. 10.1038/s41598-022-25402-z.

24. Nall, A.H., Shakhmantsir, I., Cichewicz, K., Birman, S., Hirsh, J., and Sehgal, A. (2016). Caffeine promotes wakefulness via dopamine signaling in Drosophila. Sci Rep 6, 20938. 10.1038/srep20938.

25. Wu, M.N., Ho, K., Crocker, A., Yue, Z., Koh, K., and Sehgal, A. (2009). The effects of caffeine on sleep in Drosophila require PKA activity, but not the adenosine receptor. J Neurosci 29, 11029–11037. 10.1523/JNEUROSCI.1653-09.2009.

26. Andretic, R., Kim, Y.C., Jones, F.S., Han, K.A., and Greenspan, R.J. (2008). Drosophila D1 dopamine receptor mediates caffeine-induced arousal. Proc Natl Acad Sci U S A 105, 20392–20397. 10.1073/pnas.0806776105.

27. Kayser, M.S., Mainwaring, B., Yue, Z., and Sehgal, A. (2015). Sleep deprivation suppresses aggression in Drosophila. Elife 4, e07643. 10.7554/eLife.07643.

28. Williams, M.J., Perland, E., Eriksson, M.M., Carlsson, J., Erlandsson, D., Laan, L., Mahebali, T., Potter, E., Frediksson, R., Benedict, C., and Schiöth, H.B. (2016). Recurrent Sleep Fragmentation Induces Insulin and Neuroprotective Mechanisms in Middle-Aged Flies. Front Aging Neurosci 8, 180. 10.3389/fnagi.2016.00180.

29. Silva, B., Hidalgo, S., and Campusano, J.M. (2020). Dop1R1, a type 1 dopaminergic receptor expressed in Mushroom Bodies, modulates Drosophila larval locomotion. PLoS One 15, e0229671. 10.1371/journal.pone.0229671.

30. Makos, M.A., Han, K.A., Heien, M.L., and Ewing, A.G. (2010). Using In Vivo Electrochemistry to Study the Physiological Effects of Cocaine and Other Stimulants on the Drosophila melanogaster Dopamine Transporter. ACS Chem Neurosci 1, 74–83. 10.1021/cn900017w.

31. Lim, J., Fernandez, A.I., Hinojos, S.J., Aranda, G.P., James, J., Seong, C.S., and Han, K.A. (2018). The mushroom body D1 dopamine receptor controls innate courtship drive. Genes Brain Behav 17, 158–167. 10.1111/gbb.12425.

32. Kim, Y.C., Lee, H.G., Seong, C.S., and Han, K.A. (2003). Expression of a D1 dopamine receptor dDA1/DmDOP1 in the central nervous system of Drosophila melanogaster. Gene Expr Patterns 3, 237–245. 10.1016/s1567-133x(02)00098-4.

33. Qin, H., Cressy, M., Li, W., Coravos, J.S., Izzi, S.A., and Dubnau, J. (2012). Gamma neurons mediate dopaminergic input during aversive olfactory memory formation in Drosophila. Curr Biol 22, 608–614. 10.1016/j.cub.2012.02.014.

34. Arnsten, A.F., and Pliszka, S.R. (2011). Catecholamine influences on prefrontal cortical function: relevance to treatment of attention deficit/hyperactivity disorder and related disorders. Pharmacol Biochem Behav 99, 211–216. 10.1016/j.pbb.2011.01.020.

35. Lone, S.R., Potdar, S., Srivastava, M., and Sharma, V.K. (2016). Social Experience Is Sufficient to Modulate Sleep Need of Drosophila without Increasing Wakefulness. PLoS One 11, e0150596. 10.1371/journal.pone.0150596.

36. Cheng, R.K., and Liao, R.M. (2017). Regional differences in dopamine receptor blockade affect timing impulsivity that is altered by d-amphetamine on differential reinforcement of low-rate responding (DRL) behavior in rats. Behav Brain Res 331, 177–187. 10.1016/j.bbr.2017.05.020.

37. Grant, J.E., and Chamberlain, S.R. (2018). Caffeine’s influence on gambling behavior and other types of impulsivity. Addict Behav 76, 156–160. 10.1016/j.addbeh.2017.08.007.

38. Smith, A. (2002). Effects of caffeine on human behavior. Food Chem Toxicol 40, 1243–1255. 10.1016/s0278-6915(02)00096-0.

39. Vijayraghavan, S., Major, A.J., and Everling, S. (2016). Dopamine D1 and D2 Receptors Make Dissociable Contributions to Dorsolateral Prefrontal Cortical Regulation of Rule-Guided Oculomotor Behavior. Cell Rep 16, 805–816. 10.1016/j.celrep.2016.06.031.

40. Temple, J.L., Ziegler, A.M., Graczyk, A.M., and Crandall, A. (2017). Effects of acute and chronic caffeine on risk-taking behavior in children and adolescents. J Psychopharmacol 31, 561–568. 10.1177/0269881117691568.

41. Arria, A.M., Caldeira, K.M., Kasperski, S.J., Vincent, K.B., Griffiths, R.R., and O’Grady, K.E. (2011). Energy drink consumption and increased risk for alcohol dependence. Alcohol Clin Exp Res 35, 365–375. 10.1111/j.1530-0277.2010.01352.x.

42. Pandolfo, P., Machado, N.J., Köfalvi, A., Takahashi, R.N., and Cunha, R.A. (2013). Caffeine regulates frontocorticostriatal dopamine transporter density and improves attention and cognitive deficits in an animal model of attention deficit hyperactivity disorder. Eur Neuropsychopharmacol 23, 317–328. 10.1016/j.euroneuro.2012.04.011.

43. Lazarus, M., Shen, H.Y., Cherasse, Y., Qu, W.M., Huang, Z.L., Bass, C.E., Winsky-Sommerer, R., Semba, K., Fredholm, B.B., Boison, D., et al. (2011). Arousal effect of caffeine depends on adenosine A2A receptors in the shell of the nucleus accumbens. J Neurosci 31, 10067–10075. 10.1523/JNEUROSCI.6730-10.2011.

44. Oishi, Y., and Lazarus, M. (2017). The control of sleep and wakefulness by mesolimbic dopamine systems. Neurosci Res 118, 66–73. 10.1016/j.neures.2017.04.008.

45. Guven-Ozkan, T., and Davis, R.L. (2014). Functional neuroanatomy of Drosophila olfactory memory formation. Learn Mem 21, 519–526. 10.1101/lm.034363.114.

46. de Belle, J.S., and Heisenberg, M. (1994). Associative odor learning in Drosophila abolished by chemical ablation of mushroom bodies. Science 263, 692–695. 10.1126/science.8303280.

47. Martin, J.R., Ernst, R., and Heisenberg, M. (1998). Mushroom bodies suppress locomotor activity in Drosophila melanogaster. Learn Mem 5, 179–191.

48. Pitman, J.L., McGill, J.J., Keegan, K.P., and Allada, R. (2006). A dynamic role for the mushroom bodies in promoting sleep in Drosophila. Nature 441, 753–756. 10.1038/nature04739.

49. Korshunov, K.S., Blakemore, L.J., and Trombley, P.Q. (2017). Dopamine: A Modulator of Circadian Rhythms in the Central Nervous System. Front Cell Neurosci 11, 91. 10.3389/fncel.2017.00091.

50. Le, J.Q., Ma, D., Dai, X., and Rosbash, M. (2024). Light and dopamine impact two circadian neurons to promote morning wakefulness. Curr Biol 34, 3941–3954.e3944. 10.1016/j.cub.2024.07.056.

51. Radwan, B., Liu, H., and Chaudhury, D. (2019). The role of dopamine in mood disorders and the associated changes in circadian rhythms and sleep-wake cycle. Brain Res 1713, 42–51. 10.1016/j.brainres.2018.11.031.

52. Xu, K., Zhang, Y., Shi, Y., Zhang, C., Wang, T., Lv, P., Bai, Y., and Wang, S. (2024). Circadian rhythm disruption: a potential trigger in Parkinson’s disease pathogenesis. Front Cell Neurosci 18, 1464595. 10.3389/fncel.2024.1464595.

53. Majcin Dorcikova, M., Duret, L.C., Pottié, E., and Nagoshi, E. (2023). Circadian clock disruption promotes the degeneration of dopaminergic neurons in male Drosophila. Nat Commun 14, 5908. 10.1038/s41467-023-41540-y.

54. Domaszewski, P. (2023). Gender Differences in the Frequency of Positive and Negative Effects after Acute Caffeine Consumption. Nutrients 15. 10.3390/nu15061318.

55. Temple, J.L., Bulkley, A.M., Briatico, L., and Dewey, A.M. (2009). Sex differences in reinforcing value of caffeinated beverages in adolescents. Behav Pharmacol 20, 731–741. 10.1097/FBP.0b013e328333b27c.

56. Boyer, M., Rees, S., Quinn, J., Grattan-Miscio, K., McCallum, M., and Saari, M.J. (2003). Caffeine as a performance-enhancing drug in rats: sex, dose, housing, and task considerations. Percept Mot Skills 97, 259–270. 10.2466/pms.2003.97.1.259.

57. Hughes, R.N., and Hancock, N.J. (2017). Effects of acute caffeine on anxiety-related behavior in rats chronically exposed to the drug, with some evidence of possible withdrawal-reversal. Behav Brain Res 321, 87–98. 10.1016/j.bbr.2016.12.019.

58. Turgeon, S.M., Townsend, S.E., Dixon, R.S., Hickman, E.T., and Lee, S.M. (2016). Chronic caffeine produces sexually dimorphic effects on amphetamine-induced behavior, anxiety and depressive-like behavior in adolescent rats. Pharmacol Biochem Behav 143, 26–33. 10.1016/j.pbb.2016.01.012.

59. Hughes, R.N., and Hancock, N.J. (2016). Strain-dependent effects of acute caffeine on anxiety-related behavior in PVG/c, Long-Evans and Wistar rats. Pharmacol Biochem Behav 140, 51–61. 10.1016/j.pbb.2015.11.005.

60. Machado, N.J., Ardais, A.P., Nunes, A., Szabó, E.C., Silveirinha, V., Silva, H.B., Kaster, M.P., and Cunha, R.A. (2024). Impact of Coffee Intake on Measures of Wellbeing in Mice. Nutrients 16. 10.3390/nu16172920.

61. Murillo Gonzalez, D.J., Hernandez Granados, B.A., Sabandal, P.R., and Han, K.A. (2024). Social setting interacts with hyper dopamine to boost the stimulant effect of ethanol. Addict Biol 29, e13420. 10.1111/adb.13420.

62. Shaw, P.J., Cirelli, C., Greenspan, R.J., and Tononi, G. (2000). Correlates of sleep and waking in Drosophila melanogaster. Science 287, 1834–1837. 10.1126/science.287.5459.1834.

63. Hendricks, J.C., Finn, S.M., Panckeri, K.A., Chavkin, J., Williams, J.A., Sehgal, A., and Pack, A.I. (2000). Rest in Drosophila is a sleep-like state. Neuron 25, 129–138. 10.1016/s0896-6273(00)80877-6.

64. Lin, F.J., Pierce, M.M., Sehgal, A., Wu, T., Skipper, D.C., and Chabba, R. (2010). Effect of taurine and caffeine on sleep-wake activity in Drosophila melanogaster. Nat Sci Sleep 2, 221–231. 10.2147/NSS.S13034.

65. Inan, O.T., Marcu, O., Sanchez, M.E., Bhattacharya, S., and Kovacs, G.T. (2011). A portable system for monitoring the behavioral activity of Drosophila. J Neurosci Methods 202, 45–52. 10.1016/j.jneumeth.2011.08.039.

